# Machine learning-assisted Repli-Histo labeling reveals distinct transcription-dependent constraints on chromatin motion in living cells

**DOI:** 10.64898/2026.07.05.736477

**Authors:** Katsuhiko Minami, Kako Nakazato, Sachiko Tamura, S. S. Ashwin, Kazuhiro Maeshima

**Affiliations:** Genome Dynamics Laboratory, National Institute of Genetics, ROIS, Mishima, Shizuoka 411-8540, Japan; Graduate Institute for Advanced Studies, SOKENDAI, Mishima, Shizuoka 411-8540, Japan; Department of Physics, Gandhi Institute of Technology and Management (GITAM) University, Bengaluru 561203, India

## Abstract

Genomic DNA is wrapped around core histones to form nucleosomes, which are organized in cells from euchromatin to heterochromatin with distinct genome functions. Although transcription is known to shape chromatin behavior in live cells, it remains unclear how different transcription systems shape chromatin classes and nuclear subcompartments. We developed machine learning-assisted Repli-Histo labeling to classify euchromatin and heterochromatin classes (Classes IA, IB, II, and III) and combined it with single-nucleosome imaging in live cells. Nucleosome motion was progressively constrained from euchromatin to heterochromatin. RNA polymerase II inhibition by THZ1, DRB, or α-amanitin increased nucleosome motion in euchromatic Classes IA and IB and in heterochromatin around nucleoli, but not at the nuclear periphery. In contrast, RNA polymerase I inhibition by CX-5461 selectively increased nucleosome motion in Class III heterochromatin around nucleoli. Our study reveals that Pol II and Pol I transcription shape chromatin behavior in distinct chromatin classes and nuclear subcompartments.

**Graphical Abstract:** 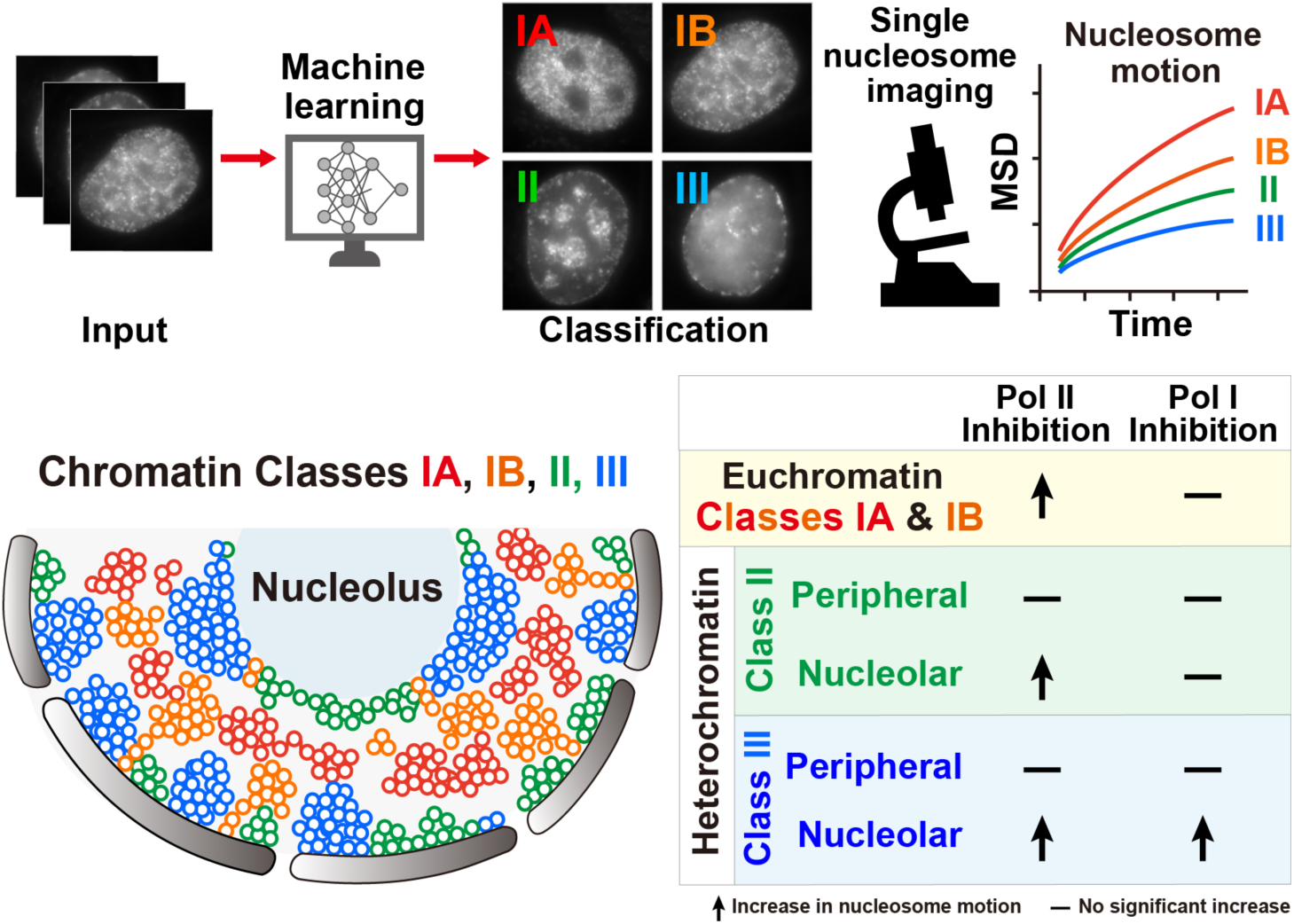

## Introduction

A long strand of DNA is wrapped around core histone octamers to form nucleosomes ^1,2^. Various lines of evidence from EM/super-resolution imaging ^3–10^ and Hi-C genomics ^11–15^ show that nucleosomes, together with other proteins and RNAs, are three-dimensionally and functionally organized into chromatin domains with distinct epigenetic marks in eukaryotic cells.

Chromatin in the cell is highly variable, ranging from euchromatin to heterochromatin, and supports various genome functions ^16–19^. Although a typical textbook view has long been that euchromatin is open and heterochromatin is closed and condensed ^20^, recent studies suggest that euchromatin domains are not fully open, but rather condensed, except for promoters and enhancers ^7,8,10,19,21–23^. Given that both euchromatin and heterochromatin form condensed domains, what is the physical difference between them? In addition to their histone modifications (e.g., active and inactive ones) and non-histone components (e.g., HP1) ^16–18^, their dynamics in live cells must be important for their functions ^24–27^.

To visualize euchromatin and heterochromatin specifically in live human cells, we recently developed replication-dependent histone labeling, termed Repli-Histo labeling ^28,29^. Repli-Histo labeling takes advantage of eukaryotic DNA replication timing: euchromatin replicates during early S phase, whereas heterochromatin replicates during late S phase (Fig. 1A). During this process, replication-dependent canonical histone variants, such as H3.2, are deposited at newly replicated nucleosomes. Based on these features, pulse labeling of newly synthesized H3.2 in early and late S phase allows specific labeling of euchromatin and heterochromatin, respectively (Fig. 1B).

**Figure 1.**
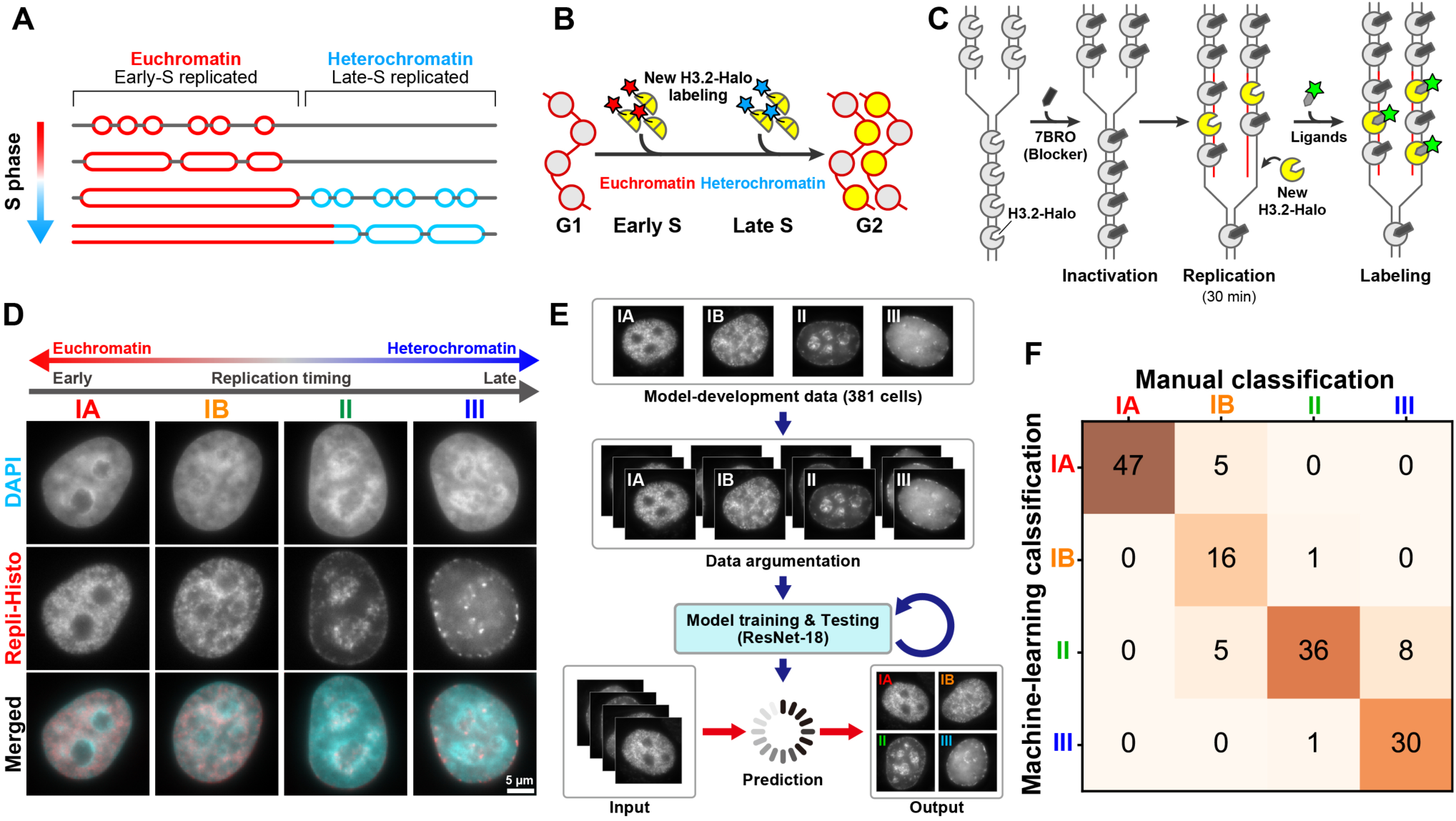
Machine-learning-assisted classification of Repli-Histo labeling patterns. **(A)** Eukaryotic cells replicate euchromatin in the early S phase, followed by heterochromatin replication in the late S phase. **(B)** The key concept of the replication-dependent histone labeling (Repli-Histo labeling). Pulse labeling of the new histones in the early and late S phases allows visualization of euchromatin and heterochromatin, respectively. **(C)** Experimental scheme of the Repli-Histo labeling. Panels (A-C) were adapted from ^28^ with minor modifications. **(D)** Representative HeLa cells with DAPI staining and Repli-Histo labeling (Classes IA, IB, II, and III from left to right). The bottom row shows the merged image (cyan, DAPI; red, Repli-Histo labeling). **(E)** Machine-learning-based classification of the Repli-Histo labeling patterns. See Methods for details. **(F)** A confusion matrix illustrating the output of machine learning predictions versus manually classified labeling patterns. Note that most of the cells fall into the diagonal categories, demonstrating that our machine-learning prediction well reproduces manual classification. The confusion matrix was generated using an independent test dataset from separate biological experiments (n = 149 cells; accuracy, 86.6%).

In Repli-Histo labeling, H3.2 fused with HaloTag (H3.2-Halo; Fig. S1A), which is distributed genome-wide, is first blocked by a non-fluorescent HaloTag ligand. Subsequently, newly synthesized H3.2-Halo molecules are incorporated into new nucleosomes and pulse-labeled with fluorescent HaloTag ligands during S phase (Fig. 1C). Based on replication timing, Repli-Histo labeling highlights four chromatin classes ranging from euchromatin to heterochromatin, namely Classes IA, IB, II, and III (Fig. 1D; Table S1). These classes correspond to historically defined replication foci ^30–32^. Early-replicating Classes IA and IB are more euchromatic, whereas later-replicating Classes II and III are more heterochromatic. These Repli-Histo classes are also aligned with Hi-C A- and B-compartments, as confirmed by pull-down and genomic sequencing ^28^. Repli-Histo labeling revealed that local nucleosome motion becomes progressively more constrained from Class IA to Class III ^28^.

How are such chromatin classes and their nucleosome motion profiles regulated and maintained in live cells? Transcription could be one of the factors that shape chromatin behavior within these classes. Previous studies, including our own, showed that active RNA Pol II (Pol II) transcription constrains chromatin motion, and that inhibition of transcription or Pol II depletion increases local chromatin motion ^5,33–39^. However, chromatin is highly heterogeneous across the genome. Euchromatin and heterochromatin differ in replication timing, histone modifications, nuclear localization, and physical properties. Therefore, it remains unclear whether transcription constrains chromatin motion relatively uniformly across the genome, or whether this constraint differs among distinct chromatin classes and nuclear subcompartments.

Here, we developed machine learning-assisted Repli-Histo labeling to objectively classify Repli-Histo patterns and combined it with single-nucleosome imaging to examine how transcription affects chromatin motion in distinct chromatin classes in living cells. Using Pol II inhibitors THZ1, DRB, and α-amanitin, we found that Pol II transcription constrains euchromatin and helps maintain the organization of heterochromatin around nucleoli. In contrast, inhibition of RNA Pol I (Pol I) transcription by CX-5461 selectively increased nucleosome motion in Class III heterochromatin around nucleoli. These results reveal distinct Pol II- and Pol I-dependent constraints on chromatin motion across chromatin classes and nuclear subcompartments in living cells.

## Results

### Machine-learning-based classification of Repli-Histo labeling patterns

Using our recently developed Repli-Histo labeling, we visualized four chromatin classes (Classes IA, IB, II, and III) in live HeLa cells based on replication timing (Figs. 1A-D)^28,29^. In our previous study, these Repli-Histo labeling patterns were classified manually. To make this classification more objective, scalable, and reproducible, we developed a machine-learning-based classifier for Repli-Histo labeling patterns (Fig. 1E). This classifier successfully categorized the four chromatin classes, with a validation accuracy of 90.5% (Fig. S1B), consistent with a related approach ^40^. The resulting model also successfully classified an independent test dataset from separate biological experiments, consistent with the manual classification (Fig. 1F; test accuracy, 86.6%). The classifier was also applicable to drug-treated cells, enabling comparisons across different experimental conditions (Fig. S1C). We hereafter refer to Repli-Histo labeling combined with this classifier as machine learning-assisted Repli-Histo labeling.

### Machine learning-assisted Repli-Histo single-nucleosome imaging maps local nucleosome motion across chromatin classes

To assess nucleosome motion in euchromatin and heterochromatin, we combined machine learning-assisted Repli-Histo labeling with single-nucleosome imaging (Fig. 2A). Two-color Repli-Histo labeling with different HaloTag ligands enabled simultaneous visualization of Repli-Histo labeling patterns and corresponding single nucleosomes: a high concentration of TMR was used to visualize the labeling patterns, whereas a very low concentration of JF646 or JFX650 ^41,42^ was used to sparsely label single nucleosomes. Using oblique illumination microscopy, which allowed us to illuminate a thin optical section within a single nucleus (Fig. 2A)^43,44^, we observed individual nucleosomes in each chromatin class as clear fluorescent dots and recorded their motion at 50 ms per frame for 200 frames, corresponding to 10 s in total (Fig. 2B; also see Movies S1-4). Each JF646 (or JFX650) dot showed single-step photobleaching, confirming that it represented a single H3.2-Halo-JF646 (or JFX650) molecule incorporated into a single nucleosome (Fig. 2C). Individual dots were then tracked using u-track software ^45^ to obtain nucleosome trajectories (Fig. 2D). The position determination accuracy of fluorescent H3.2-Halo-JF646 dots was 10.9 nm (Fig. S2A).

**Figure 2.**
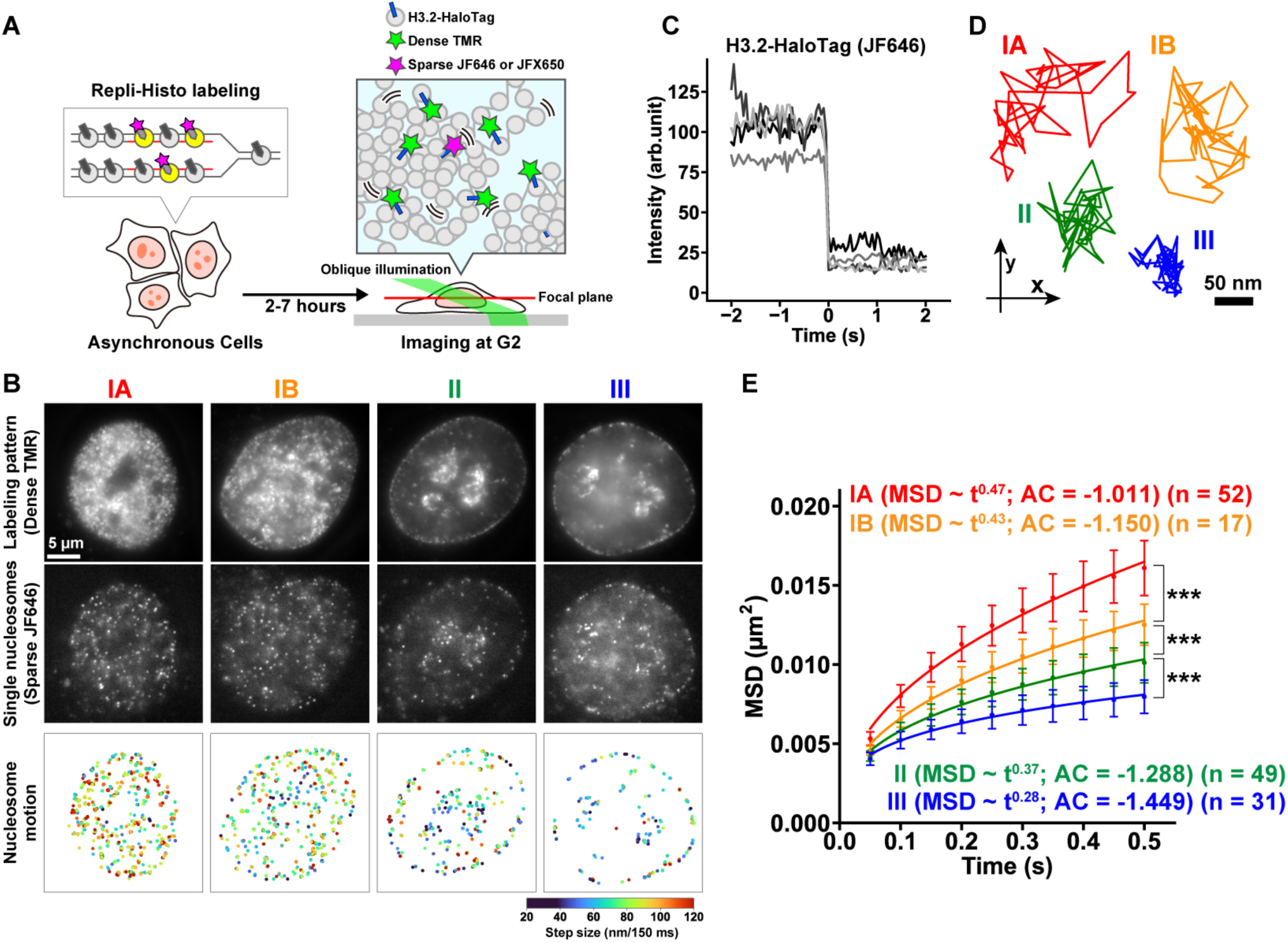
Machine learning-assisted Repli-Histo single-nucleosome imaging maps local nucleosome motion across chromatin classes. (A) Schematic of the single-nucleosome imaging. (Left) The S phase HeLa cells were labeled with Repli-Histo labeling. (Right) The Repli-Histo-labeled cells were observed at the following G2 phase using an oblique illumination microscopy system. A small fraction of H3.2-Halo was labeled with JF646 or JFX650 (magenta) for single-nucleosome imaging, and the rest was densely labeled with TMR (green) to visualize labeling patterns. The panel was adapted from ^28^ with modifications. (B) HeLa cells with two-color Repli-Histo labeling of the Classes IA, IB, II, and III regions. From top to bottom: labeling patterns (dense TMR), single nucleosomes (sparse JF646), heatmaps of the single-nucleosome displacement. (C) Single-step photobleaching of five representative nucleosomes (H3.2-Halo-JF646). The horizontal axis shows the time before and after photobleaching. (D) Representative trajectories of tracked single nucleosomes in Classes IA (red), IB (orange), II (green), and III (blue) chromatin in live HeLa cells. (E) MSD plots (± SD among cells) of single nucleosomes in IA (red, n = 52), IB (orange, n = 17), II (green, n = 49), and III (blue, n = 31) in living HeLa cells. Note that chromatin labeling patterns were determined and sorted by our machine-learning workflow (Fig. 1E-F). MSD exponents and the AC values are from Fig. S2B and E.

As reported in our previous study, nucleosome motion became progressively more constrained from Class IA to Class III (Fig. 2E). Nucleosomes in heterochromatin classes, Classes II and III, were more constrained than those in euchromatin classes, Classes IA and IB. The MSD exponent α also decreased along this chromatin class axis: 0.47 for Class IA, 0.43 for Class IB, 0.37 for Class II, and 0.28 for Class III (Fig. S2B). We examined the moving-angle (motion-vector) distribution of individual nucleosomes (Fig. S2C) and calculated the asymmetry coefficient (AC) values (Fig. S2D) ^46,47^. Nucleosomes in more heterochromatic chromatin classes have more pulling back force (i.e., a smaller AC value; Fig. S2E). These results are highly consistent with our previous findings ^28^, further supporting the reliability of our machine learning-based chromatin class classification (Fig. 1E-F).

### RNA Pol II inhibition by THZ1 increases local nucleosome motion

To ask how transcription affects chromatin behavior in each class, we treated cells with THZ1, a covalent inhibitor of CDK7, a kinase involved in RNA Pol II (Pol II) CTD phosphorylation and transcription initiation ^48^. Previous studies, including our own, have shown that transcription constrains chromatin motion in living cells ^5,33–39,49^. Immunostaining for Pol II CTD serine 5 phosphorylation (Ser5P), a marker associated with transcription initiation, showed a significant reduction after THZ1 treatment (Fig. 3A-B). Consistently, THZ1 markedly suppressed global RNA synthesis, as validated by reduced 5-ethynyl uridine (EU) incorporation (Fig. 3C-D). In agreement with previous single-nucleosome imaging studies ^33^, THZ1 increased genome-wide nucleosome motion, as measured by sparse H3.2-Halo single-nucleosome imaging (Fig. S3A).

**Figure 3.**
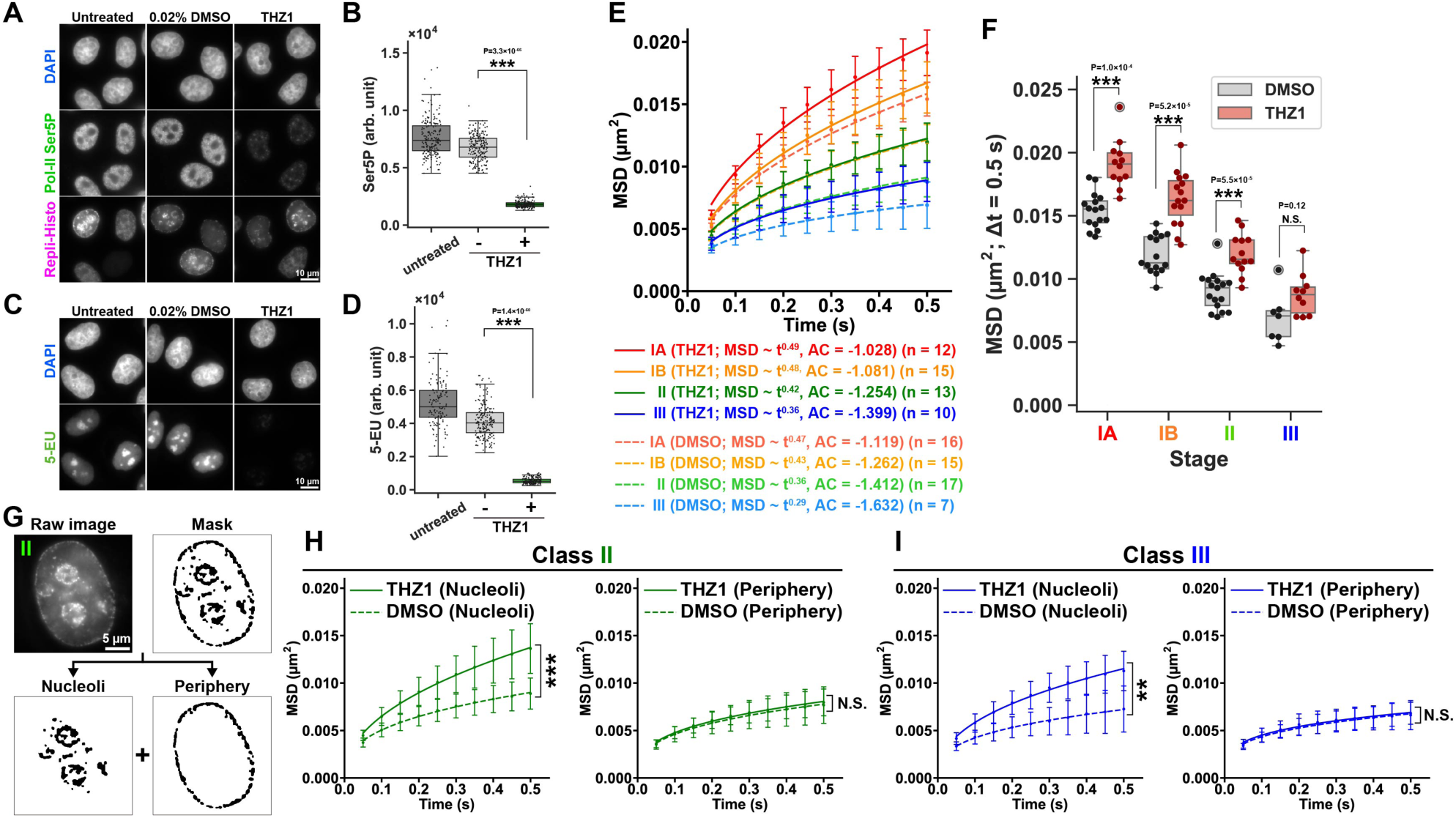
RNA Pol II inhibition increases local nucleosome motion across chromatin classes. **(A)** RNA Pol II inhibitor THZ1 significantly reduces active Pol-II marker Ser5P levels monitored by immunofluorescence. As controls, untreated cells and DMSO-treated cells were also shown. From top to bottom: DAPI staining, anti-RNA polymerase II Ser5-phosphorylation (Pol-II Ser5P) immunofluorescence, and Repli-Histo labeling. **(B)** The quantification of Pol-II Ser5P levels in the untreated (n = 201 cells), DMSO-treated (n = 213 cells), and THZ1-treated cells (n = 184 cells). **(C)** Transcription levels after THZ1 treatment were also validated by 5-EU incorporation into nascent RNA. The incorporated 5-EU was detected using Alexa Fluor 488 picolyl azide via click chemistry. **(D)** The quantification of 5-EU intensity in the untreated (n = 134 cells), DMSO-treated (n = 193 cells), and THZ1-treated cells (n = 169 cells). **(E)** MSD plots (± SD among cells) of single nucleosomes in IA (DMSO, n = 16; THZ1, n = 12), IB (DMSO, n = 15; THZ1, n = 15), II (DMSO, n = 17; THZ1, n = 13), and III (DMSO, n = 7; THZ1, n = 10) in DMSO (dashed lines) or THZ1 (solid lines) treated cells from 0.05 to 0.5 s. MSD exponents and AC values are from Fig. S3B. **(F)** MSD at 0.5 s in each cell from (E). ***: *P* = 1.0 × 10^-4^ (IA; DMSO vs THZ1), *P* = 5.2 × 10^-5^ (IB; DMSO versus THZ1), *P* = 5.5 × 10^-5^ (II; DMSO versus THZ1). N.S.: *P* = 0.12 (III; DMSO versus THZ1). Data are tested by the two-sided Kolmogorov-Smirnov test. **(G)** Classes II and III chromatin are subcategorized into nucleolus-associated regions (nucleoli) and peri-nuclear regions (periphery) and are analyzed separately. **(H)** MSD plots (± SD among cells) of nucleoli- and periphery-associated single nucleosomes in the Class II chromatin. ***: *P* = 5.5 × 10^-5^ (nucleoli; DMSO versus THZ1), N.S.: *P* = 0.58 (periphery; DMSO versus THZ1) by the two-sided Kolmogorov-Smirnov test. **(I)** MSD plots (± SD among cells) of nucleoli- and periphery-associated single nucleosomes in the Class III chromatin. **: *P* = 0.0017 (nucleoli; DMSO versus THZ1), N.S.: *P* = 0.63 (periphery; DMSO versus THZ1) by the two-sided Kolmogorov-Smirnov test.

We then treated Repli-Histo-labeled cells with THZ1 for 2 h and performed machine learning-assisted Repli-Histo single-nucleosome imaging. Local nucleosome motion in the four chromatin classes was observed during the subsequent G2 phase (Fig. 2A; Movies S1-S4). Local nucleosome motion in Classes IA and IB was markedly increased after Pol II transcription inhibition (Figs. 3E-F, S3B; Movies S5-S8).

### THZ1 distinctly affects heterochromatin nucleosome motion at the nuclear periphery and around nucleoli

Interestingly, THZ1 treatment significantly increased nucleosome motion in Class II heterochromatin (Fig. 3E-F). Nucleosome motion in Class III heterochromatin also showed a slight increase, although the difference was not statistically significant (Fig. 3E-F). To examine this observation in more detail, we subdivided Classes II and III heterochromatin into regions at the nuclear periphery and around nucleoli (Fig. 3G), and analyzed their nucleosome motion separately. As reported previously, regions within the same chromatin class showed similar nucleosome motion under control conditions (DMSO; Fig. 3H-I) ^28,29^. In contrast to euchromatin, local nucleosome motion in perinuclear heterochromatin of Classes II and III, which largely corresponds to lamina-associated domains (LADs) ^50^, was not affected by THZ1 treatment (right; Fig. 3H-I). By contrast, THZ1 significantly increased nucleosome motion in heterochromatin around nucleoli, which is enriched in nucleolus-associated domains (NADs) ^51^ (left; Fig. 3H-I). These results suggest that Pol II inhibition differentially affects heterochromatin motion at the nuclear periphery and around nucleoli.

### Pol II transcription constrains euchromatin and helps maintain heterochromatin around nucleoli

To test the generality of these results, we performed similar experiments using two other Pol II inhibitors, DRB and α-amanitin (α-AM). DRB inhibits CDK9 kinase in the P-TEFb complex, which phosphorylates Ser2 in the C-terminal domain of the largest subunit of Pol II, RPB1 ^52,53^. DRB also dissociates the elongating Pol II complex from chromatin ^54^. α-AM inhibits Pol II more directly by binding near its catalytic active site ^55^ and induces degradation of RPB1 ^56^. Immunostaining for RPB1 Ser5P showed a significant reduction after DRB or α-AM treatment (Figs. 4A-B and S4C), validating that these inhibitors effectively inhibited Pol II transcription.

**Figure 4.**
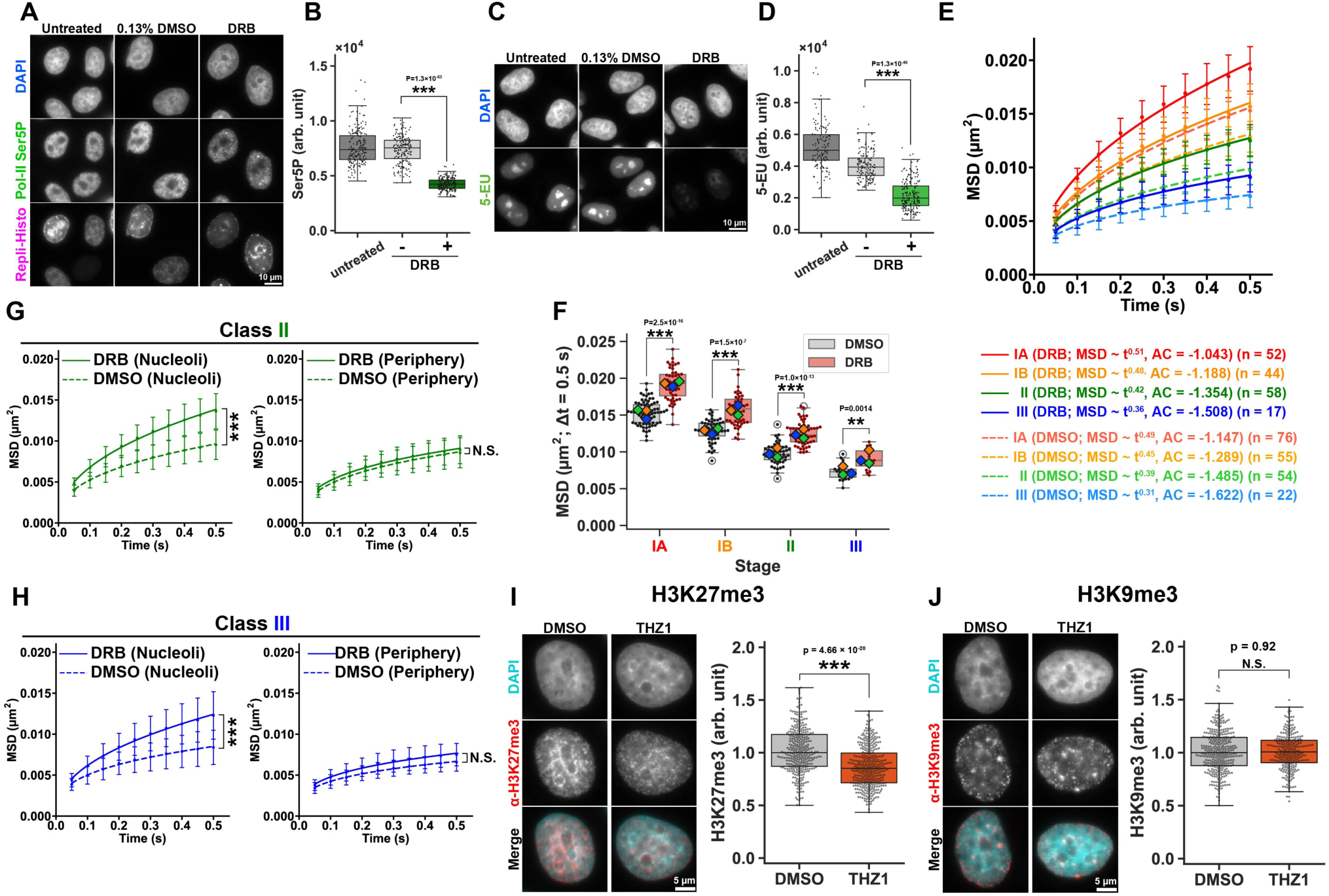
Pol II transcription constrains euchromatin and helps maintain heterochromatin around nucleoli. **(A)** DRB significantly reduces active Pol-II marker Ser5P levels monitored by immunofluorescence. As controls, untreated cells (reproduced from Fig. 3A) and DMSO-treated cells were also shown. From top to bottom: DAPI staining, anti-RNA polymerase II Ser5-phosphorylation (Pol-II Ser5P) immunofluorescence, and Repli-Histo labeling. **(B)** The quantification of Pol-II Ser5P levels in the untreated (n = 201 cells), DMSO-treated (n = 188 cells), and DRB-treated cells (n = 205 cells). **(C)** Transcription level after DRB treatment was also validated by 5-EU incorporation into nascent RNA. The incorporated 5-EU was detected using Alexa Fluor 488 picolyl azide via click chemistry. The untreated control was reproduced from Fig. 3C. **(D)** The quantification of 5-EU intensity in the untreated (n = 134 cells), DMSO-treated (n = 173 cells), and DRB-treated cells (n = 211 cells). **(E)** MSD plots (± SD among cells) of single nucleosomes in IA (DMSO, n = 76; DRB, n = 52), IB (DMSO, n = 55; DRB, n = 44), II (DMSO, n = 54; DRB, n = 58), and III (DMSO, n = 22; DRB, n = 17) in DMSO (dashed lines) or DRB (solid lines) treated HeLa cells from 0.05 to 0.5 s. MSD exponents and AC values are from Fig. S4B. **(F)** MSD at 0.5 s in each cell from (E). Large diamonds show mean values in each replicate. ***: *P* = 2.5 × 10^-16^ (IA; DMSO vs DRB), *P* = 1.5 × 10^-7^ (IB; DMSO versus DRB), *P* = 1.0 × 10^-13^ (II; DMSO versus DRB). **: *P* = 0.0014 (III; DMSO versus DRB). Data are tested by the two-sided Kolmogorov-Smirnov test. **(G)** MSD plots (± SD among cells) of nucleoli- and periphery-associated single nucleosomes in the Class II chromatin. ***: *P* = 2.6 × 10^-15^ (nucleoli; DMSO versus DRB), N.S.: *P* = 0.70 (periphery; DMSO versus DRB) by the two-sided Kolmogorov-Smirnov test. **(H)** MSD plots (± SD among cells) of nucleoli- and periphery-associated single nucleosomes in the Class III chromatin. ***: *P* = 1.1 × 10^-5^ (nucleoli; DMSO versus DRB), N.S.: *P* = 0.075 (periphery; DMSO versus DRB) by the two-sided Kolmogorov-Smirnov test. **(I)** THZ1 treatment significantly decreases H3K27me3 levels in HeLa cells, monitored by immunofluorescence. (Left) Representative images. From top to bottom: DAPI staining, anti-H3K27me3 immunofluorescence, and merged images (cyan, DAPI; red, H3K27me3). (Right) The quantification of H3K27me3 levels from the left. **(J)** THZ1 treatment did not significantly decrease H3K9me3 levels. (Left) Representative images. From top to bottom: DAPI staining, anti-H3K9me3 immunofluorescence, and merged images (cyan, DAPI; red, H3K9me3). (Right) The quantification of H3K9me3 levels from the left.

Consistent with the results using THZ1, DRB treatment significantly increased nucleosome motion in Classes IA and IB euchromatin (Fig. 4E-F, Fig. S4A-B). In addition, DRB treatment increased nucleosome motion in Classes II and III heterochromatin. To examine this effect in more detail, we separately analyzed nucleosome motion at the nuclear periphery and around nucleoli in Classes II and III (Fig. 4G-H). We again found that DRB treatment significantly increased nucleosome motion only in heterochromatin around nucleoli, whereas heterochromatin at the nuclear periphery did not change. Notably, we obtained very similar findings with α-AM treatment (Fig. S4C-G).

Furthermore, we observed consistent results using human RPE-1 cells expressing H3.2-HaloTag (Fig. S5A) ^28^. THZ1 treatment increased nucleosome motion in Classes IA and IB euchromatin and in Class II heterochromatin around nucleoli, while nucleolus-associated Class III heterochromatin showed the same tendency (Fig. S5B-D). The differential chromatin constraint by Pol II transcription seems to be a general feature in human cells.

We next asked why heterochromatin motion around nucleoli increased after Pol II inhibition. To gain insight into this question, we examined the heterochromatin-associated histone marks H3K9me3 and H3K27me3 after THZ1 treatment. THZ1 significantly reduced H3K27me3 signals, whereas H3K9me3 signals remained largely unchanged (Fig. 4I-J). This selective reduction of H3K27me3 suggests that Pol II inhibition perturbs H3K27me3-associated facultative heterochromatin, including that around nucleoli, which may contribute to the increased nucleosome motion in these regions. Taken together, our findings indicate that Pol II transcription constrains Classes IA and IB euchromatin and is also involved in maintaining heterochromatin around nucleoli. Once Pol II transcription was inhibited, heterochromatin around nucleoli was altered, and its nucleosome motion increased.

### RNA Pol I transcription selectively constrains Class III heterochromatin around nucleoli

Since we found that Pol II transcription affects not only euchromatin but also heterochromatin behavior, we next asked how RNA Pol I (Pol I) transcription affects chromatin behavior. To address this question, we treated Repli-Histo-labeled cells with CX-5461, an RNA Pol I inhibitor ^57,58^. CX-5461 treatment severely suppressed RNA synthesis in nucleoli, as validated by reduced EU incorporation (Fig. 5A-B). In addition, immunostaining signals of fibrillarin, a marker of the dense fibrillar component of nucleoli, were decreased after CX-5461 treatment (Fig. 5A, C), suggesting that CX-5461 perturbed nucleolar organization.

**Figure 5.**
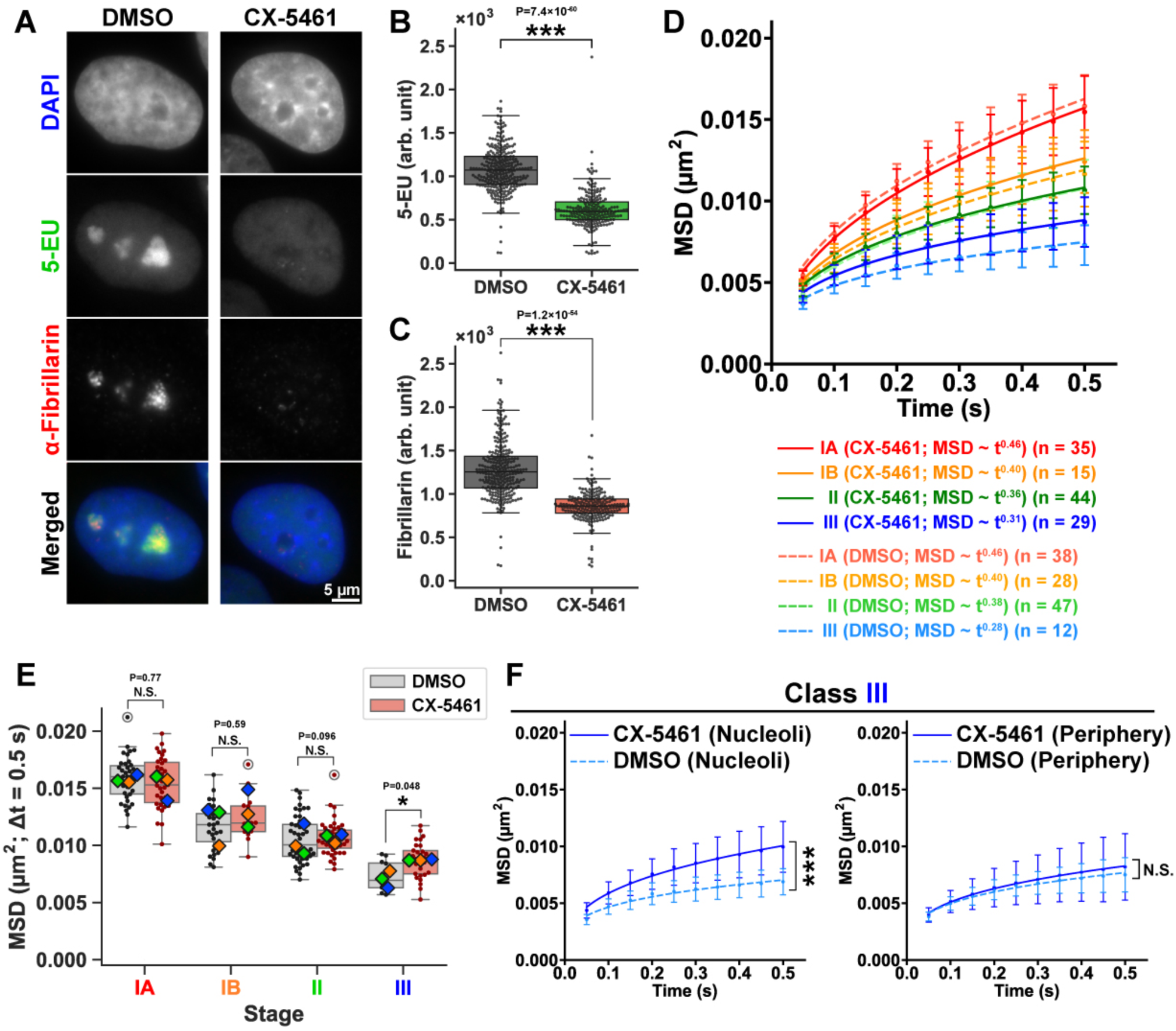
RNA Pol I locally constrains nucleolus-associated heterochromatin but not other chromatin regions. **(A)** Pol-I inhibitor CX-5461 treatment reduces nascent RNA transcription in the nucleoli of HeLa cells, as validated by 5-EU labeling via click chemistry. Perturbation of nucleolar organization was validated by anti-fibrillarin immunofluorescence. From top to bottom: DAPI staining, 5-EU, anti-fibrillarin immunofluorescence, merged images (blue, DAPI; green, 5-EU; red, fibrillarin). **(B and C)** The quantification of 5-EU and fibrillarin levels from (A). **(D)** MSD plots (± SD among cells) of single nucleosomes in IA (DMSO, n = 38; CX-5461, n = 35), IB (DMSO, n = 28; CX-5461, n = 15), II (DMSO, n = 47; CX-5461, n = 44), and III (DMSO, n = 12; CX-5461, n = 29) in DMSO (dashed lines) or CX-5461 (solid lines) treated cells from 0.05 to 0.5 s. MSD exponents are from Fig. S6A. **(E)** MSD at 0.5 s in each cell from (D). Large diamonds show mean values in each replicate. N.S.: *P* = 0.77 (IA; DMSO vs CX-5461), *P* = 0.59 (IB; DMSO versus CX-5461), *P* = 0.096 (II; DMSO versus CX-5461). *: *P* = 0.048 (III; DMSO versus CX-5461). Data are tested by the two-sided Kolmogorov-Smirnov test. **(F)** MSD plots (± SD among cells) of nucleoli- and periphery-associated single nucleosomes in Class III chromatin. ***: *P* = 4.3 × 10^-5^ (nucleoli; DMSO versus CX-5461), N.S.: *P* = 0.71 (periphery; DMSO versus CX-5461) by the two-sided Kolmogorov-Smirnov test.

We then performed machine learning-assisted Repli-Histo single-nucleosome imaging after 2 h of CX-5461 treatment. In contrast to Pol II inhibition, CX-5461 did not significantly increase nucleosome motion in Classes IA, IB, or II, but caused a modest increase in Class III chromatin (Figs. 5D-E, S6A). Further subdivision of Class III regions revealed that this response was mainly observed in nucleolar regions, whereas peripheral regions showed no significant change (Fig. 5F). Consistently, the overlap of Class III heterochromatin foci with NPM1-positive regions was significantly reduced after CX-5461 treatment (Fig. S6B-D). These results indicate that Pol I transcription locally constrains Class III heterochromatin around nucleoli.

Thus, Pol II and Pol I inhibition produced distinct chromatin-class and nuclear-subcompartment responses: Pol II inhibition affected euchromatin and nucleolar heterochromatin, whereas Pol I inhibition selectively affected Class III heterochromatin around nucleoli.

## Discussion

In this study, we developed machine learning-assisted Repli-Histo labeling and combined it with single-nucleosome imaging to examine nucleosome motion in distinct chromatin classes in living cells. Our previous Repli-Histo study showed that local nucleosome motion becomes progressively constrained from early-replicating euchromatin (Classes IA and IB) to late-replicating heterochromatin (Classes II and III)^28,29^. Here, by replacing manual classification with a machine-learning-based classifier, we objectively categorized Repli-Histo labeling patterns (Fig. 1E-F) and compared chromatin behavior across different perturbation conditions. This approach revealed distinct transcription-dependent physical constraints on nucleosome motions across the genome. Pol II inhibition increased nucleosome motion in euchromatin classes and heterochromatin around nucleoli, whereas Pol I inhibition selectively increased nucleosome motion in Class III heterochromatin around nucleoli. These findings extend Repli-Histo labeling from a method for mapping physical properties of euchromatin and heterochromatin to a platform for dissecting chromatin-class- and nuclear-subcompartment-specific regulatory mechanisms.

How does Pol II transcription constrain Classes IA and IB euchromatin, which are transcriptionally active regions? Although the mechanism remains unclear, there are several possibilities. The first is that chromatin is globally and physically stabilized by loose connections through active Pol II ^33^, which is compatible with classical transcription factory models ^59,60^. Recent studies have shown that active Pol II, Mediator, and other transcription factors form dynamic clusters/droplets, presumably through phase separation or related processes ^61–66^. These transcription condensates are thought to associate with and physically constrain transcribing chromatin regions, thereby supporting efficient transcription and transcriptional regulation. Indeed, BRD4-NUT transcriptional condensates constrained neighboring nucleosome motion via acetylated histones ^67^. Another theoretical study proposed that active mechanical forces generated by transcribing RNA Pol II can suppress chromatin motion through transient ordering of gene-rich chromatin ^68^. Finally, transcribed RNA molecules themselves may also contribute to constraining chromatin. Nuclear RNAs are negatively charged long polymers and, together with RNA-binding proteins, can form RNA-rich meshworks or gel-like structures that influence chromatin architecture ^69^. Such RNA-dependent nuclear structures may contribute to transcription-dependent constraints on chromatin motion.

We also found that Pol II transcription helps maintain heterochromatin around nucleoli, but not heterochromatin at the nuclear periphery. Notably, distinct drugs (THZ1, DRB, and α-AM), which have different primary targets in the Pol II transcription machinery (CDK7, CDK9, and the Pol II catalytic subunit, respectively), produced similar responses, validating our results. Thus, late-replicating heterochromatin is not physically uniform: Heterochromatin at the nuclear periphery is associated with LADs and may be constrained by lamina-dependent tethering and H3K9me3-rich constitutive heterochromatin mechanisms. In contrast, heterochromatin around nucleoli is enriched in NADs and is embedded in a nucleolar/perinucleolar environment. In our experiments, THZ1 reduced H3K27me3 signals while H3K9me3 signals remained largely unchanged. This suggests that Pol II inhibition preferentially perturbs H3K27me3-associated facultative heterochromatin, including that around nucleoli, rather than globally disrupting H3K9me3-rich heterochromatin. Therefore, the increase in nucleosome motion around nucleoli after Pol II inhibition may reflect reduced physical constraints on facultative heterochromatin associated with nucleolar/perinucleolar organization. Indeed, nucleolar Pol II itself may contribute to nucleolar organization ^70^.

Pol I inhibition produced a distinct response from Pol II inhibition. Pol I inhibitor CX-5461 suppressed ribosomal RNA (rRNA) synthesis ^57^ and perturbed NPM1-positive nucleolar organization ^58,71^, but it did not significantly increase nucleosome motion in Classes IA, IB, and II. Instead, its effect was mainly observed in Class III heterochromatin around nucleoli. Since rRNA transcribed by Pol I maintains nucleolar organization, rRNA may constrain late-replicating Class III heterochromatin associated with nucleoli. While Pol I inhibition induces the condensation and segregation of fibrillar center (FC) components to the periphery, forming spherical “nucleolar caps” ^58^, Pol II inhibition drives FC components to decondense into thread-like “nucleolar necklaces” ^72^. Indeed, we observed that Class III heterochromatin around nucleoli partially dissociates from the NPM1-positive body, representing the granular component, the outermost compartment of nucleoli, after CX-5461 treatment (Fig. S6B-D), further supporting a physical constraint mediated by rRNA-dependent nucleolar organization.

In summary, machine learning-assisted Repli-Histo labeling revealed that transcription-dependent chromatin constraints differ among chromatin classes and nuclear subcompartments in living cells. It should be emphasized that all perturbations were performed after Repli-Histo labeling, so that euchromatin/heterochromatin labeling had already been established before the perturbations, ensuring reliable comparisons across conditions. We conclude that Pol II transcription constrains euchromatic chromatin and helps maintain heterochromatin around nucleoli, whereas Pol I transcription locally constrains Class III heterochromatin around nucleoli. These findings support a model in which chromatin behavior is regulated not only by euchromatin/heterochromatin identity, but also by transcriptional systems and nuclear subcompartmental context. Thus, Repli-Histo labeling provides a powerful framework for dissecting how genome functions are coupled to the physical behavior of chromatin in living cells.

## Methods

### Cell lines

HeLaS3 ^73^ and RPE-1 (CRL-4000; ATCC) cells expressing endogenous human H3.2 tagged with HaloTag were established as described previously ^28^ and were cultured at 37 °C in 5% CO_2_ in Dulbecco’s Modified Eagle’s medium (DMEM) (D5796- 500ML; Sigma-Aldrich) supplemented with 10% fetal bovine serum (F7524; Sigma-Aldrich).

### Repli-Histo labeling

The replication-dependent histone labeling (Repli-Histo labeling) was performed as described previously ^28^. First, asynchronous cells expressing an endogenous H3.2-HaloTag were incubated with the medium supplemented with 10 µM 7-bromo-1-heptanol (7BRO, B1852; Tokyo Chemical Industry) ^74^ for 60 min at 37 °C in 5% CO_2_. The cells were then washed three times with medium without 7BRO. The cells were then exposed to the medium supplemented with 100 nM HaloTag TMR ligand for 30 min immediately after washing out 7BRO. Subsequently, the cells were washed three times with 1× HBSS and treated with 10 µM 7BRO for 30 min to block any remaining unreacted HaloTag sites. The cells were then treated with chemical inhibitors as described below. Finally, the cells were fixed with 1.85% formaldehyde (064–00406; Wako) for 15 min, permeabilized with 0.5% Triton X-100 (T-9284; Sigma Aldrich) for 5 min, and stained with 0.5 µg/mL DAPI (10236276001; Roche) for 5 min before being embedded in PPDI [20 mM HEPES (pH 7.4), 1 mM MgCl_2_, 100 mM KCl, 78% glycerol, and 1 mg/mL paraphenylene diamine (PPDI, 695106-1G; Sigma-Aldrich)].

### Chemical treatment

For transcription inhibition, the Repli-Histo-labeled cells were treated with transcription inhibitors, 100 μM 5,6-dichlorobenzimidazole 1-β-D-ribofuranoside (DRB) (D1916-10MG; Sigma-Aldrich) for 2 h, 100 μg/mL α-amanitin (α-AM; 010-22961; Fujifilm Wako) for 2 h, and 1 μM THZ1 ^48^ (CS-3168; ChemScene) for 2 h. For RNA polymerase I inhibition, cells were treated with 1 μM CX-5461 (CS-0568; ChemScene) for 2 h.

### Machine-learning-based image classification

To characterize the replication timing of the labeled regions, the labeled patterns (historically called replication foci) of the Repli-Histo labeling were categorized into the four classes (IA, IB, II, and III) based on previous studies ^31^.

For automated classification of Repli-Histo labeling patterns into the four classes (IA, IB, II, III), we developed a deep learning pipeline based on the ResNet18 architecture ^75^. The pipeline consists of three main components: data processing, image preprocessing, and model architecture (Fig. 1E). We first performed Repli-Histo labeling on asynchronous HeLa cells as described above and collected 381 nuclear images for model development. The dataset was split into training (265 images, 70%) and validation (116 images, 30%) subsets. Data were processed in batches of 32 samples using PyTorch’s DataLoader. Image preprocessing utilized torchvision transforms. Images were converted to 8-bit grayscale (num output channels = 1) and resized to 224 × 224 pixels. The pixel values of each image were rescaled to a pixel-value range of 0–255. Training data augmentation included random horizontal flips, random rotations (± 10 degrees), and random resized crops (scale factor 0.8-1.0). The processed images were converted to tensors and normalized with mean and standard deviation values of 0.5. The model was based on a pre-trained ResNet18 architecture with modifications. The first convolutional layer was adapted for single-channel grayscale input (7 × 7 kernel, 64 output channels, stride = 2, padding = 3). The network included dropout (*P* = 0.7), batch normalization, and a modified final fully connected layer for four-class output. The training was performed using the Adam optimizer ^76^ with a learning rate of 0.0001 and cross-entropy loss for 50 epochs. The trained model was validated using the validation subset (validation accuracy: 90.5%). We finally applied the model to an independent test dataset from separate biological experiments (149 cells; test accuracy, 86.6%). We confirmed that the predicted labeling patterns were consistent with those that were manually classified (Fig. 1F).

### Western blotting

Cells were lysed in Laemmli sample buffer ^77^ supplemented with 10% 2-mercaptoethanol (133-1457; Wako) and incubated at 95 °C for 5 min to denature proteins. The cell lysates, equivalent to 1 × 10^5^ cells per well, were subjected to SDS–polyacrylamide gel electrophoresis (12.5% for histone detection). For western blotting, the fractionated proteins in the gel were transferred to a polyvinylidene difluoride (PVDF) membrane (IPVH00010; Millipore) by a semi-dry blotter (BE-320; BIO CRAFT). After blocking with 5% skim milk (190-12865; Fujifilm Wako), the membrane-bound proteins were probed by the mouse anti-H3.1/H3.2 (1:1000; CEC-006; Cosmo Bio) or the mouse anti–HaloTag (G9281; Promega, 1:1,000) followed by the anti-mouse IgG (170-6516; Bio-Rad, 1:5,000) horseradish peroxidase-conjugated goat antibody. Bands were detected by chemiluminescence reactions (WBKLS0100; Millipore) and images were acquired with the EZ-Capture MG (AE-9300H-CSP; ATTO).

### Indirect immunofluorescence

Immunostaining was performed as described previously ^78^, and all processes were performed at room temperature. Cells were fixed with 1.85% FA in PBS for 15 min and then treated with 50 mM glycine in HMK [20 mM HEPES (pH 7.5) with 1 mM MgCl_2_ and 100 mM KCl] for 5 min and permeabilized with 0.5% Triton X-100 in HMK for 5 min. After washing twice with HMK for 5 min, the cells were incubated with 10% normal goat serum (NGS, 143-06561; Wako) in HMK for 30 min. The cells were incubated with the diluted primary antibody, mouse anti–phosphorylated Ser5 of RNAPII (gift from Prof. H. Kimura, 1:1000), mouse anti-H3K27me3 (gift from Prof. H. Kimura, 1:1000), anti fibrillarin (ab4566; Abcam, 1:500), anti NPM1 (B0556; Sigma, 1:1000) in 1% NGS in HMK for 1 h. For H3K9me3 detection, cells were incubated with mouse anti-H3K9me3 (gift from Prof. H. Kimura, 1:1000), diluted in Can Get Signal^®^ immunostain Immunoreaction Enhancer Solution (NKB601; TOYOBO) to improve immunoreaction specificity. After being washed with HMK four times, the cells were incubated with the diluted secondary antibody, goat anti–rabbit IgG Alexa Fluor 488 (1:500, A-11006; Thermo Fisher Scientific), or goat anti–mouse IgG Alexa Fluor 647 (1:500, A-21235; Thermo Fisher Scientific) in 1% NGS in HMK (or the Enhancer Solution) for 1 h followed by a wash with HMK four times. For DNA staining in fixed cells, 0.5 μg/mL DAPI was added to the cells for 5 min followed by washing with HMK. The stained cells were embedded in PPDI.

### EU labeling and detection

EU incorporation was performed by using Click-iT RNA Alexa Fluor 488 Imaging Kit (C10329; Thermo Fisher Scientific) in combination with Alexa Fluor 488 picolyl azide (C10637; Thermo Fisher Scientific). Cells were incubated with 500 μM EU for 1 h during the period of chemical treatment. The cells were then fixed with 1.85% FA for 15 min and permeabilized with 0.5% Triton X-100 for 5 min. Incorporated EU was labeled by click reaction according to the manufacturer’s instructions, followed by quenching with the Click-iT reaction rinse buffer (C10329; Thermo Fisher Scientific). The cells were finally stained with 0.5 μg/mL DAPI for 5 min and embedded in PPDI.

### Fluorescent microscopy on fixed samples

Image stacks of fixed cells were acquired by using a Delta Vision Ultra (GE Healthcare) with an Olympus UPLXAPO 60× (NA 1.42) objective. Optical sections at a thickness of 0.2 µm were imaged. The best-focused optical section near the middle of each nucleus was extracted after correcting the chromatic aberration. Image analysis was performed using Fiji/ImageJ.

To quantify the intranuclear signal intensity of Repli-Histo labeling, DAPI-stained nuclear regions were segmented based on Huang’s fuzzy thresholding method with Fiji/ImageJ, and the total nuclear pixel intensities (arb. unit) and mean nuclear intensities (arb. unit) for DAPI and all other channels were measured. The sample numbers for each experiment are shown in the corresponding figure legends.

### Single-nucleosome imaging microscopy

For single-nucleosome imaging, established cell lines were cultured on poly-L-lysine (P1524-500MG; Sigma-Aldrich) coated glass-based dishes (3970-035; Iwaki). H3.2-Halo molecules were fluorescently labeled with 40 pM HaloTag TMR ligand for 20 min at 37 °C in 5% CO_2_. The cells were washed with Hank’s balanced salt solution (1× HBSS, H1387; Sigma-Aldrich) three times and then incubated in a phenol red-free DMEM (21063–029; Thermo Fisher Scientific) for over 2 h before live-cell imaging.

For the Repli-Histo labeling, asynchronous cells were first treated with 10 µM 7BRO as described above. The cells were then washed three times with medium without 7BRO, followed by a 30-minute pulse labeling with a mixture of 2 nM HaloTag TMR ligand and 300 pM HaloTag JFX650 (or 5 nM HaloTag JF646) ligand. After washing three times with a medium, cells were once more treated with 10 µM 7BRO for 30 minutes to block any remaining unreacted HaloTag sites. The cells were finally subjected to live-cell imaging after treatment with the chemical inhibitors.

Single-nucleosome imaging was performed as described previously ^28^. To maintain cell culture conditions (37 °C, 5% CO_2_, and humidity) under the microscope, a live-cell chamber and a digital gas mixer (STXG-WSKMX-SET; Tokai Hit) were used. Single nucleosomes were observed by using an inverted Nikon Eclipse Ti2 microscope, an ILE 400 laser combiner (Andor) with 100-mW 561-nm and 140-mW 637-nm laser systems, and a sCMOS ORCA-Fusion BT camera (C15440-20UP, Hamamatsu Photonics). Fluorescently labeled histones in living cells were excited by the 561/637-nm laser through an objective lens (100× PlanApo TIRF, NA 1.49; Nikon) and detected at 582-626/676-786 nm. An oblique illumination system with the TIRF unit (Nikon) was used to excite the labeled histone molecules within a limited thin area in the cell nucleus and reduce background noise. Sequential image frames were acquired using NIS Elements software (AR v5.30.03 64-bit, Nikon) at 50 ms per frame under continuous illumination.

To determine the position determination accuracy of the single-nucleosome dots, we performed single-nucleosome imaging on chemically fixed cells. The fluorescently labeled cells grown on poly–L–lysine coated glass-based dishes were fixed with 2% formaldehyde in 1×PBS at 37 °C for 15 min and washed with 1×PBS three times. The following single-nucleosome imaging was performed as described above.

### Single-nucleosome tracking analysis

The methods for image processing, single-molecule tracking, and single-nucleosome movement analysis were described previously ^5^. The background noise signals in the acquired sequential TIFF images were subtracted with the rolling ball background subtraction (rolling ball radius: 50 pixels) using Fiji/ImageJ. The nuclear regions in the images were manually extracted. Following this step, the diffraction-limited fluorescent dots in each image were fitted to a Gaussian function to obtain the center of the distribution, and its trajectory was tracked with u-track (MATLAB package ^45^). To ascertain the position determination accuracy of the H3.2-HaloTag nucleosomes in FA-fixed cells, distributions of nucleosome displacements from the centroids of the trajectories in the x-and y-planes (n = 10 molecules) were fitted to Gaussian functions. The calculated position determination accuracy in each experiment was described in Fig. S2A.

For single-nucleosome movement analysis, the displacement and MSD of the fluorescent dots were calculated based on their trajectory using a Python script. The originally calculated MSD was in 2D. To obtain the 3D value, the 2D value was multiplied by 1.5 (4 to 6 *Dt*). The calculated MSDs were fitted to a sub-diffusive model *MSD*(*t*) = 6*D* · *t^α^*, where α is an anomalous diffusion exponent (0 < α < 1). Statistical analyses of the single-nucleosome MSD were performed using Python.

### Segmentation of the Class II/III chromatin regions

To extract the single nucleosome trajectories colocalized with late-S replication foci (Fig. 3G), we segmented the late-S replicated chromatin regions as described previously ^28^. First, 10-frame (0.5 s) averaged TMR images were processed with an ImageJ band-pass filter function with the filtering of large structures down to 30 pixels and small structures up to 5 pixels. The filtered images were then binarized using Otsu’s method. The 2-pixel-dilated binary masks were defined as the late-S replication foci region. The detected single nucleosome dots located inside the foci regions were extracted based on the centroids of their trajectories (i.e., time average of xy-coordinates) as described previously ^28,67^. The categorized trajectories were used in subsequent analyses.

### Colocalization of the Class III chromatin and nucleoli

We evaluated the association of Class III chromatin with nucleoli by immunofluorescence against NPM1, a marker of the outermost nucleolar compartment (granular component). To extract the NPM1 regions, we first processed the raw images with an ImageJ top-hat filter (radius = 20 pixels), followed by denoising with a Gaussian blur filter (sigma = 1.5 pixels). The filtered images were binarized using the max-entropy method and then eroded by 2 pixels to cancel the expansion caused by the Gaussian blur filter. The resulting binary regions (including holes) were defined as NPM1-positive nucleoli. The Class III chromatin was labeled with Repli-Histo labeling. The acquired images were processed using an ImageJ band-pass filter, filtering large structures down to 12 pixels and small structures up to 1.8 pixels. The filtered images were then binarized using Otsu’s method. The resulting binary masks were defined as the Class III chromatin regions.

To examine the colocalization of Class III regions and NPM1 bodies, we merged their binarized images and calculated the fraction of nucleoplasmic Class III pixels that overlapped with NPM1-positive regions on a pixel-by-pixel basis. The intersection coefficient was calculated as the number of overlapping pixels divided by the total number of nucleoplasmic Class III pixels. The analysis was performed using the Python Scikit-image library [https://scikit-image.org/docs/0.25.x/auto_examples/applications/plot_colocalization_metrics.html#colocalization-metrics].

### Angle distribution analysis

Angle distribution analysis was performed as described in ^46^. For the tracked consecutive points {(***x***_0_, ***y***_0_), (***x***_1_, ***y***_1_), ⋯ , (***x****_n_*, ***y****_n_*), ⋯ } of a single nucleosome on the xy-plane, we converted the data into a set of displacement vectors, Δ***r****_n_* = (*x_n_*_+1_ – *x_n_*, *y_n_*_+1_ – *y_n_*)^t^. Then, we calculated the angle between two vectors Δ***r****_n_* and Δ***r****_n_*_+1_. We carried out this procedure for all the points of each trajectory in our experiments. We plotted the normalized polar histogram by our Python program ^79^. The angle distribution was normalized by 2π, and the values correspond to the probability density.

## Acknowledgments

We are grateful to Dr. K. M. Marshall for critical reading of this manuscript. We thank Mr. J. Hernandez for initial results and Dr. S. Ide for technical assistance. We thank Dr. H. Kimura for providing H3K9me3, H3K27me3, and RNA Pol II antibodies. Finally, we thank all members of the Maeshima laboratory for valuable discussions and continuous support.

## Author Contributions

K.Minami and K.Maeshima designed the research. K.Minami and K. N. performed the experiments including cell generation, imaging, and analyses. S.S.A., K.Minami, and K.N. developed machine-learning methods for Repli-Histo labeling. S.T. and K.Maeshima performed some biochemical experiments and some figure illustrations. K.Minami and K.Maeshima wrote the manuscript with input from all authors.

## Data and Materials Availability

All data needed to evaluate the conclusions in the paper are present in the paper and/or the Supplementary Materials. The scripts for track-sorting and angle-distribution analysis are available at https://zenodo.org/records/12672197. The scripts for machine-learning are available at https://doi.org/10.5281/zenodo.21186176.

## Supplementary Movie Legends

**Movie S1.** A HeLa cell with Class IA Repli-Histo labeling, treated with 0.02% DMSO. (Left) The Class IA Repli-Histo labeling pattern with dense TMR labeling. (Right) A movie (50 ms/frame) of the corresponding single nucleosomes labeled with JFX650. Images were acquired with the sCMOS ORCA-Fusion BT (Hamamatsu Photonics). Note that clear and well-separated JFX650 dots show single-step photobleaching profiles (Fig. 2C), suggesting that each dot represents a single H3.2-Halo-JFX650 molecule in a single nucleosome. Scale bar: 5 µm.

**Movie S2.** A HeLa cell with Class IB Repli-Histo labeling, treated with 0.02% DMSO. (Left) The Class IB Repli-Histo labeling pattern with dense TMR labeling. (Right) A movie (50 ms/frame) of the corresponding single nucleosomes labeled with JFX650. Scale bar: 5 µm.

**Movie S3.** A HeLa cell with Class II Repli-Histo labeling, treated with 0.02% DMSO. (Left) The Class II Repli-Histo labeling pattern with dense TMR labeling. (Right) A movie (50 ms/frame) of the corresponding single nucleosomes labeled with JFX650. Scale bar: 5 µm.

**Movie S4.** A HeLa cell with Class III Repli-Histo labeling, treated with 0.02% DMSO. (Left) The Class III Repli-Histo labeling pattern with dense TMR labeling. (Right) A movie (50 ms/frame) of the corresponding single nucleosomes labeled with JFX650. Rapidly diffusing dots are free H3.2-Halo, and we focused only on the behavior of H3.2-Halo that is stably incorporated into nucleosomes. Scale bar: 5 µm.

**Movie S5.** A HeLa cell with Class IA Repli-Histo labeling, treated with 1 µM THZ1. (Left) The Class IA Repli-Histo labeling pattern with dense TMR labeling. (Right) A movie (50 ms/frame) of the corresponding single nucleosomes labeled with JFX650. Scale bar: 5 µm.

**Movie S6.** A HeLa cell with Class IB Repli-Histo labeling, treated with 1 µM THZ1. (Left) The Class IB Repli-Histo labeling pattern with dense TMR labeling. (Right) A movie (50 ms/frame) of the corresponding single nucleosomes labeled with JFX650. Scale bar: 5 µm.

**Movie S7.** A HeLa cell with Class II Repli-Histo labeling, treated with 1 µM THZ1. (Left) The Class II Repli-Histo labeling pattern with dense TMR labeling. (Right) A movie (50 ms/frame) of the corresponding single nucleosomes labeled with JFX650. Scale bar: 5 µm.

**Movie S8.** A HeLa cell with Class III Repli-Histo labeling, treated with 1 µM THZ1. (Left) The Class III Repli-Histo labeling pattern with dense TMR labeling. (Right) A movie (50 ms/frame) of the corresponding single nucleosomes labeled with JFX650. Rapidly diffusing dots are free H3.2-Halo, and we focused only on the behavior of H3.2-Halo that is stably incorporated into nucleosomes. Scale bar: 5 µm.

---

**Figure S1.**
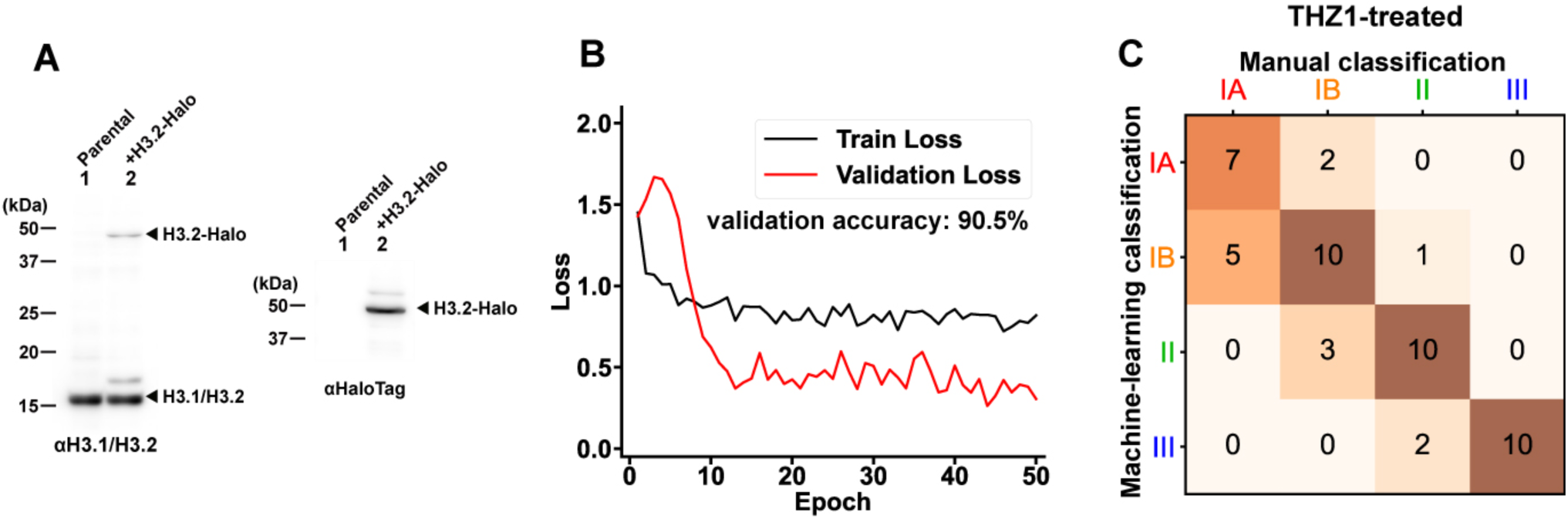
Validation of the machine-learning-assisted Repli-Histo labeling pattern classification. **(A)** Western blots of histone H3.2-HaloTag in lysates of HeLa cells (lane 2) using an anti-histone H3.1/H3.2 antibody (left) and an anti-HaloTag antibody (right). **(B)** Training and validation losses of a representative model over epochs during the training process. **(C)** A confusion matrix illustrating the output of machine learning predictions versus manual classification of THZ1-treated HeLa cells.

**Figure S2.**
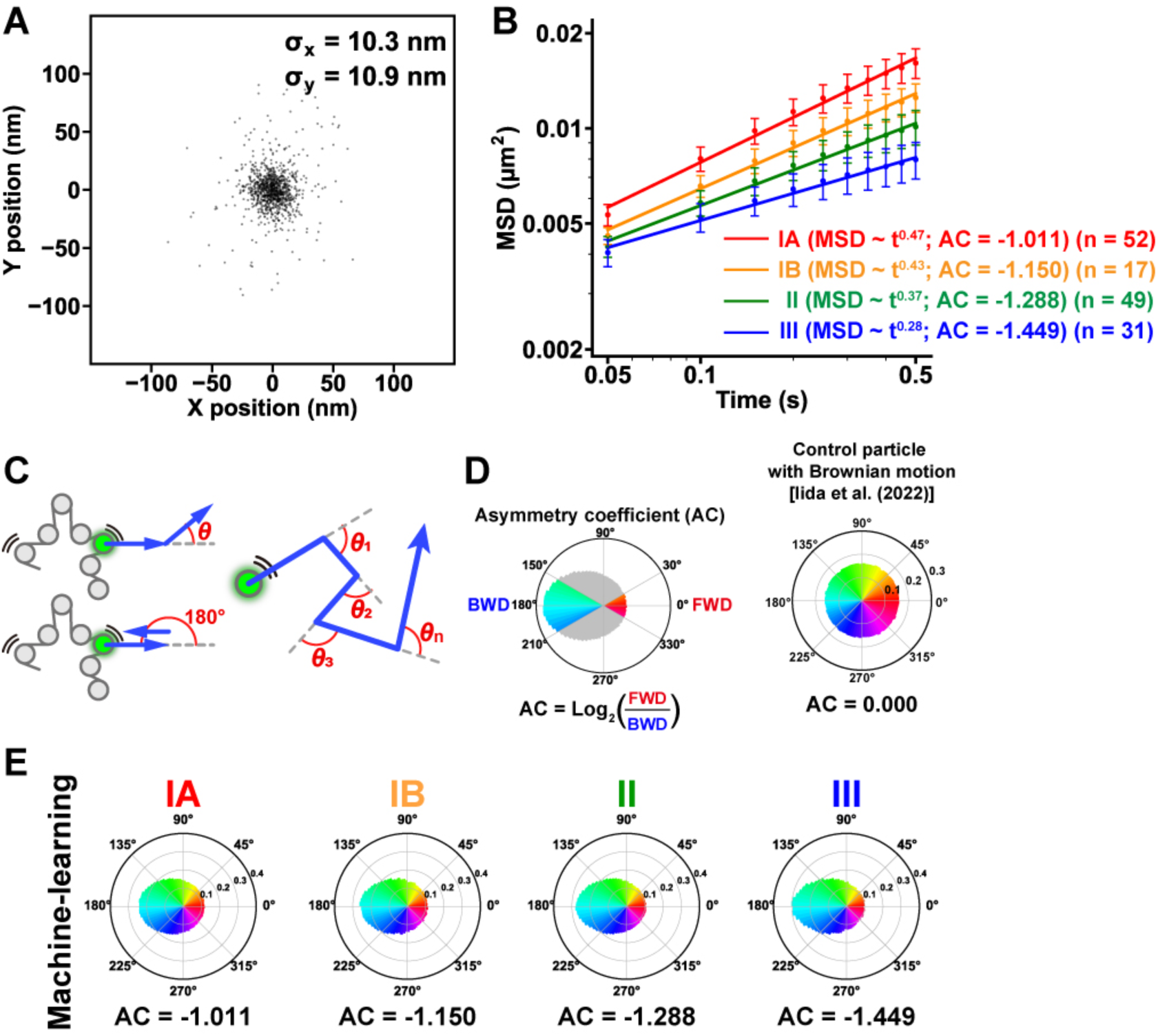
Local nucleosome movement in the four classes of chromatin regions. **(A)** The position determination accuracy of JF646-labeled nucleosomes. Distribution of nucleosome displacements from the centroids in the XY-plane, n = 10 nucleosomes in FA-fixed HeLa cells. SDx and SDy of fitted Gaussian functions were 10.3 nm and 10.9 nm, respectively. **(B)** The log-log plot of MSDs from the plot of Fig. 2E. The plots were fitted linearly. The anomalous exponent values calculated from the fitted lines are shown. **(C)** Schematic for angle-distribution analysis. **(D)** (Left) Schematic for the asymmetry coefficient (AC). See Methods for details. (Right) Angle distribution from a particle with Brownian motion [reproduced from ^46^]. **(E)** Motion angle distributions of nucleosomes in Classes IA, IB, II, and III in living HeLa cells (data from Fig. 2E). Their AC values are shown at the bottom.

**Figure S3.**
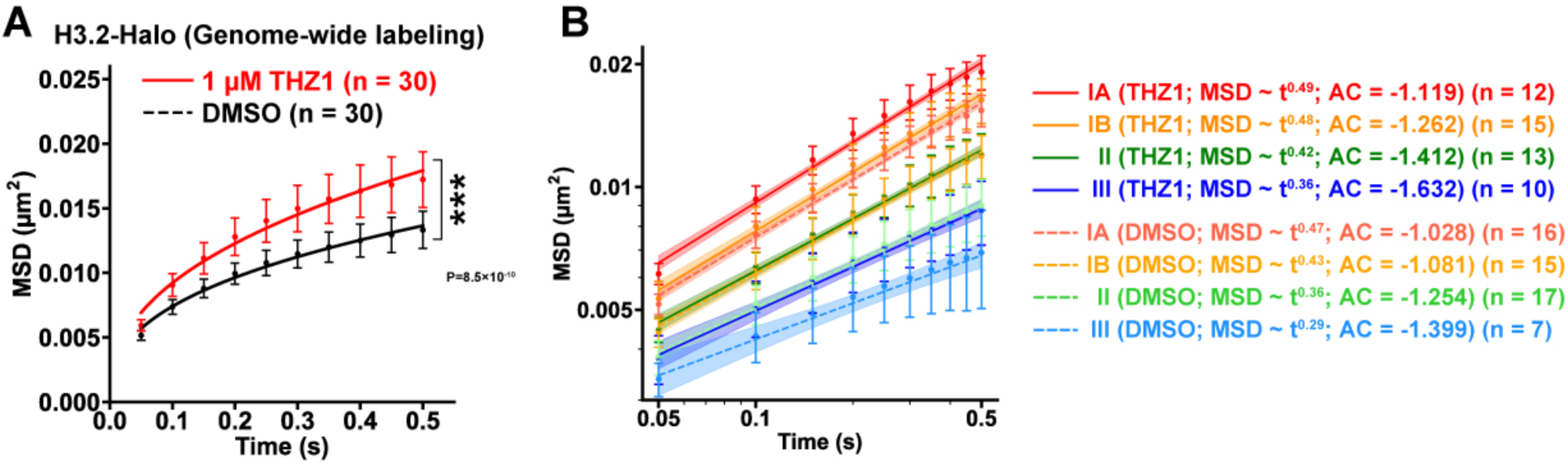
THZ1 treatment increases local nucleosome motion in HeLa cells. **(A)** MSD plots (±SD among cells) of average nucleosome motion in the HeLa cells treated with 1 µM THZ1 (red) or DMSO (black). For each condition, n = 30 cells were analyzed. **(B)** The log-log plot of MSDs from Fig. 3E. The plots were fitted linearly. The anomalous exponent values calculated from the fitted lines and the AC values from the motion angle distribution analysis are also shown.

**Figure S4.**
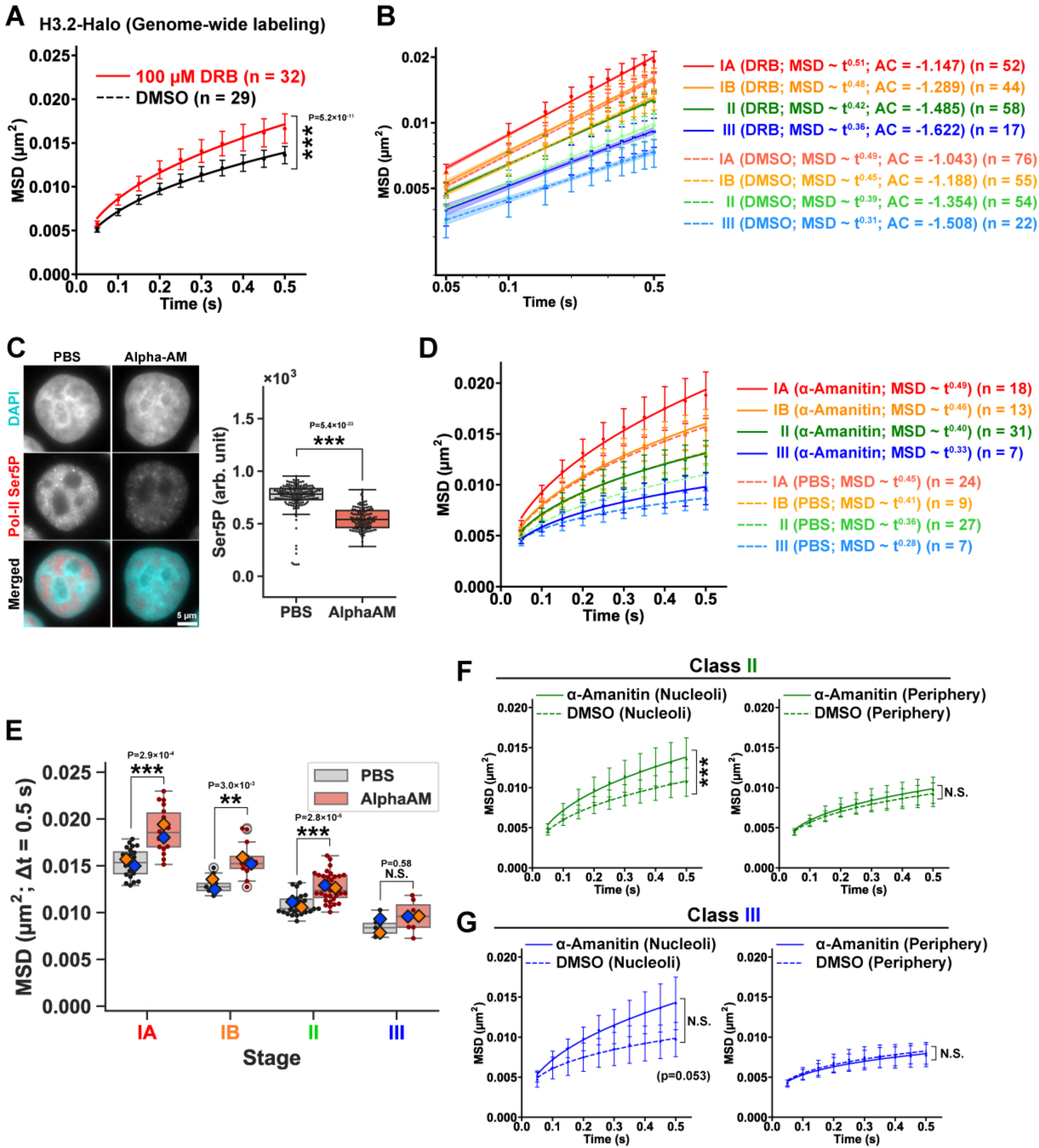
Various RNA Pol II inhibitors generally increase local nucleosome motion. **(A)** MSD plots (±SD among cells) of average nucleosome motion in the HeLa cells treated with 100 µM DRB (red; n = 32) or DMSO (black; n = 29). **(B)** The log-log plot of MSDs from Fig. 4E. The plots were fitted linearly. The anomalous exponent values calculated from the fitted lines and the AC values from the motion angle distribution analysis are also shown. **(C)** (Left) The Pol-II inhibitor α- amanitin (α-AM) significantly reduces the Pol-II Ser5P level, as monitored by immunofluorescence. As controls, 1×PBS-treated cells were also shown. From top to bottom: DAPI staining, immunostaining of Pol-II Ser5P, and merged image. (Right) The quantification of Pol-II Ser5P levels from the left. **(D)** MSD plots (± SD among cells) of single nucleosomes in IA (PBS, n = 24; α-AM, n = 18), IB (PBS, n = 9; α-AM, n = 13), II (PBS, n = 27; α-AM, n = 31), and III (PBS, n = 7; α-AM, n = 7) in PBS (dashed lines) or α-AM(solid lines) treated cells from 0.05 to 0.5 s. MSD exponents and AC values are also shown. **(E)** MSD at 0.5 s in each cell from (D). Large diamonds show mean values in each replicate. ***: *P* = 2.9 × 10^-4^ (IA; PBS vs α-AM), *P* = 3.0 × 10^-3^ (IB; PBS versus α-AM), *P* = 2.8 × 10^-5^ (II; PBS versus α-AM). N.S.: *P* = 0.58 (III; PBS versus α-AM). Data are tested by the two-sided Kolmogorov-Smirnov test. **(F)** MSD plots (± SD among cells) of nucleoli- and periphery-associated single nucleosomes in Class II chromatin. ***: *P* = 5.0 × 10^-5^ (nucleoli; PBS versus α-AM), N.S.: *P* = 0.26 (periphery; PBS versus α-AM) by the two-sided Kolmogorov-Smirnov test. **(G)** MSD plots (± SD among cells) of nucleoli- and periphery-associated single nucleosomes in Class III chromatin. N.S.: *P* = 0.053 (nucleoli; PBS versus α-AM), *P* = 0.99 (periphery; PBS versus α-AM) by the two-sided Kolmogorov-Smirnov test.

**Figure S5.**
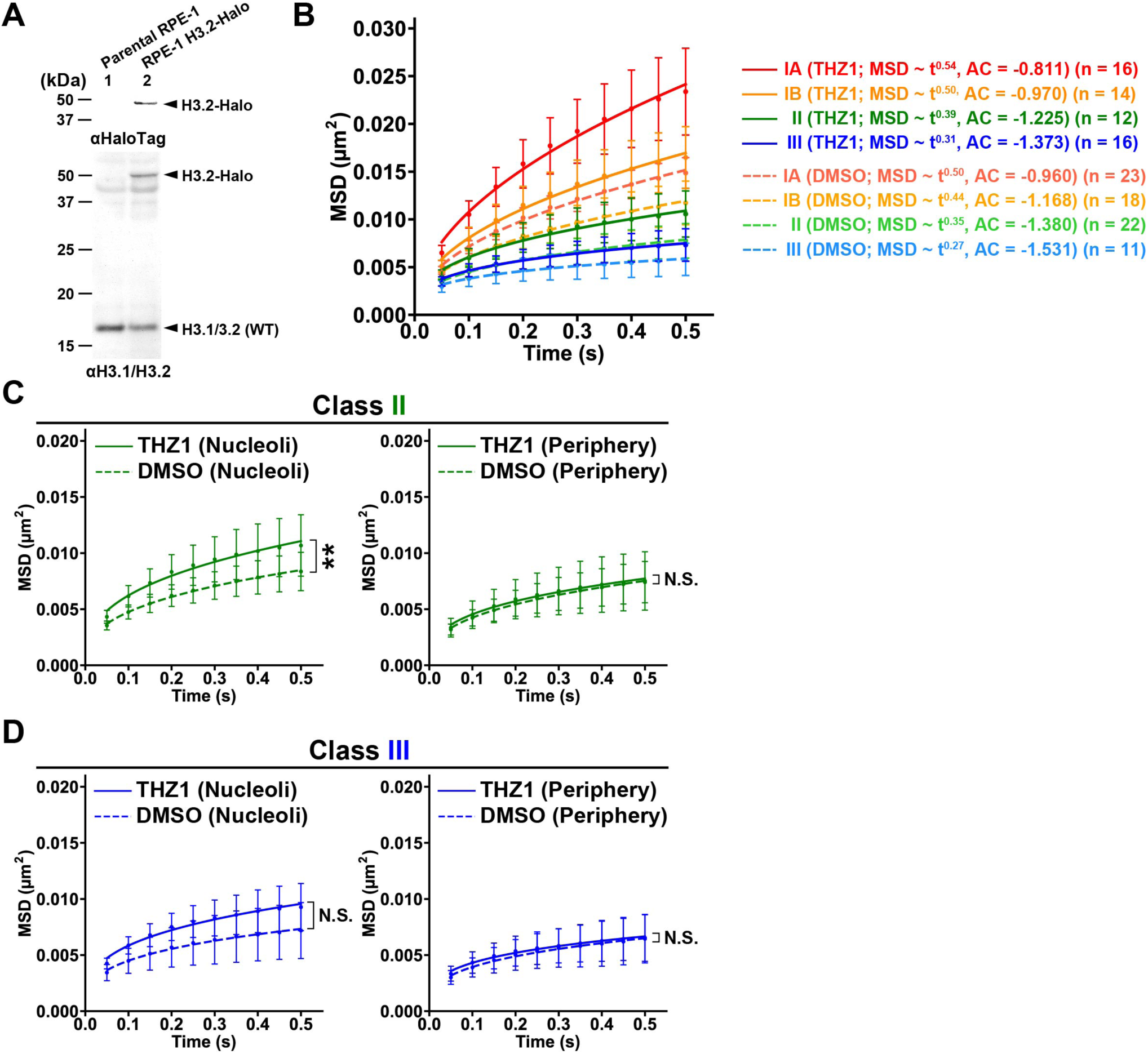
THZ1 treatment increases local nucleosome motion in RPE-1 cells. **(A)** Western blots of histone H3.2-HaloTag in lysates of RPE-1 cells (lane 2) using an anti-HaloTag antibody (top) and an anti-histone H3.1/H3.2 antibody (bottom). **(B)** MSD plots (± SD among cells) of single nucleosomes in IA (DMSO, n = 23; THZ1, n = 16), IB (DMSO, n = 18; THZ1, n = 14), II (DMSO, n = 22; THZ1, n = 12), and III (DMSO, n = 11; THZ1, n = 16) in DMSO (dashed lines) or THZ1 (solid lines) treated RPE-1 cells from 0.05 to 0.5 s. MSD exponents and AC values are also shown. **(C)** MSD plots (± SD among cells) of nucleoli- and periphery-associated single nucleosomes in Class II chromatin of RPE-1 cells. **: *P* = 5.8 × 10^-3^ (nucleoli; DMSO versus THZ1), N.S.: *P* = 0.49 (periphery; DMSO versus THZ1) by the two-sided Kolmogorov-Smirnov test. **(D)** MSD plots (± SD among cells) of nucleoli- and periphery-associated single nucleosomes in Class III chromatin of RPE-1 cells. N.S.: *P* = 0.16 (nucleoli; DMSO versus THZ1), N.S.: *P* = 0.96 (periphery; DMSO versus THZ1) by the two-sided Kolmogorov-Smirnov test. Notably, because Class III cells were labeled during late S phase, fewer Class III cells remained in G2 phase at the time of imaging, likely reflecting the relatively short G2 phase of RPE-1 cells ^80^. Consequently, the sample size for this class was smaller.

**Figure S6.**
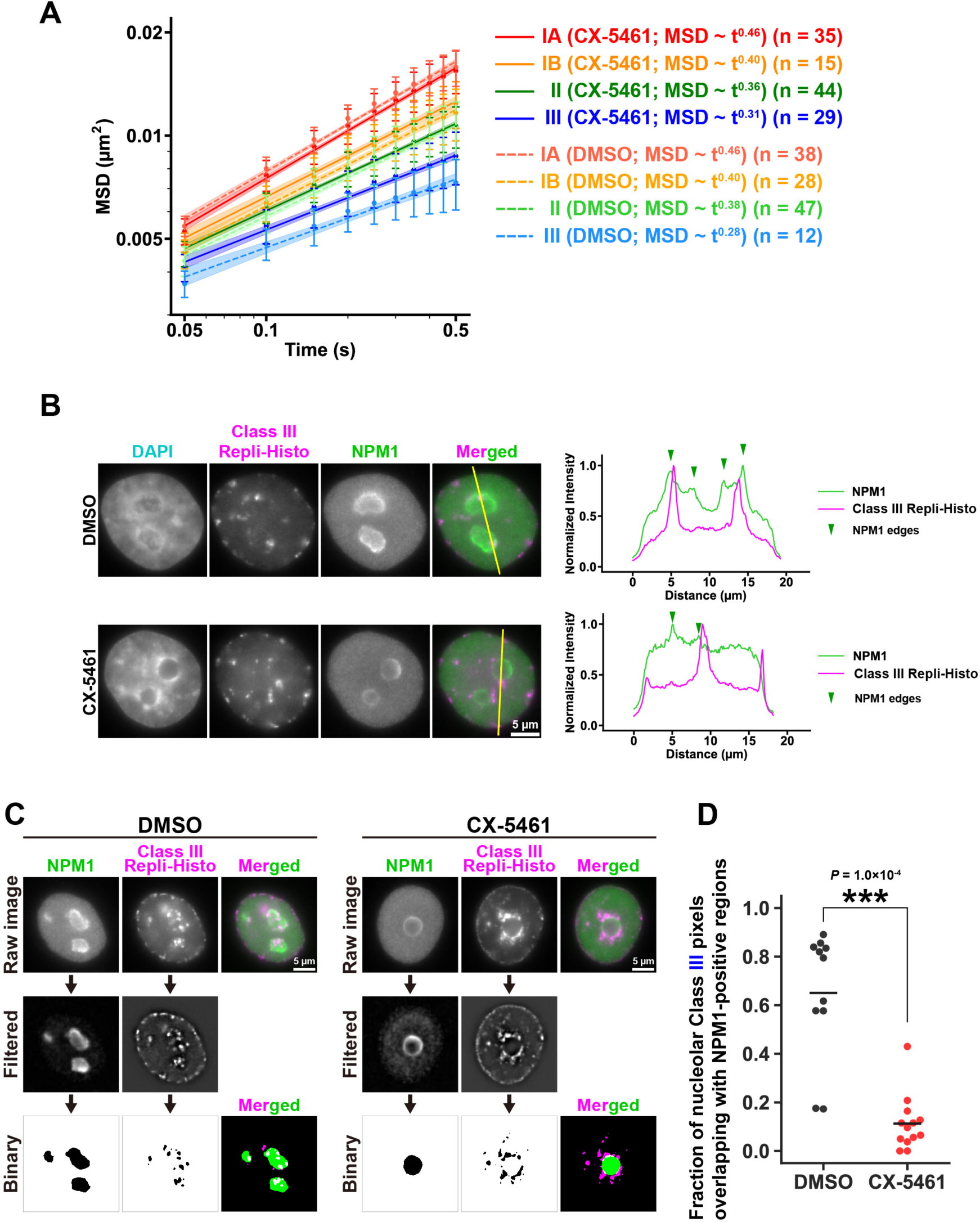
CX-5461 treatment perturbs nucleolus-associating Class III heterochromatin organization. **(A)** The log-log plot of MSDs from the plot of Fig. 5D. The plots were fitted linearly. The anomalous exponent values calculated from the fitted lines are shown. **(B)** CX-5461 treatment partially disrupts the nucleolar association of Class III heterochromatin foci. Representative Class III-labeled HeLa nuclei are shown as an example. From left to right: DAPI staining, Repli-Histo labeling, anti-NPM1 immunofluorescence (nucleolus GC component), and merged images (magenta, Repli-Histo labeling; green, NPM1). (right) The intensity line profiles of NPM1 (green) and Repli-Histo labeling (magenta) across the yellow lines in the left panels. Note that the nucleolus-associated Repli-Histo foci are partially relocated from within to outside the nucleoli after CX-5461 treatment. **(C)** The colocalization of nucleolar-associated Class III regions and NPM1 bodies was assessed by the overlap of their binarized images (intersection coefficient; see Methods for details). **(D)** The quantified colocalization coefficient (intersection coefficient). The fraction of nucleolar-associated Class III pixels that overlapped with NPM1-positive regions was calculated on a pixel-by-pixel basis. n = 11 cells (DMSO) and n = 13 cells (CX-5461) were analyzed. Black bars show the mean among cells. ***: *P* = 1.0 × 10^-4^ by two-sided Mann-Whitney *U* test.

**Table S1:**
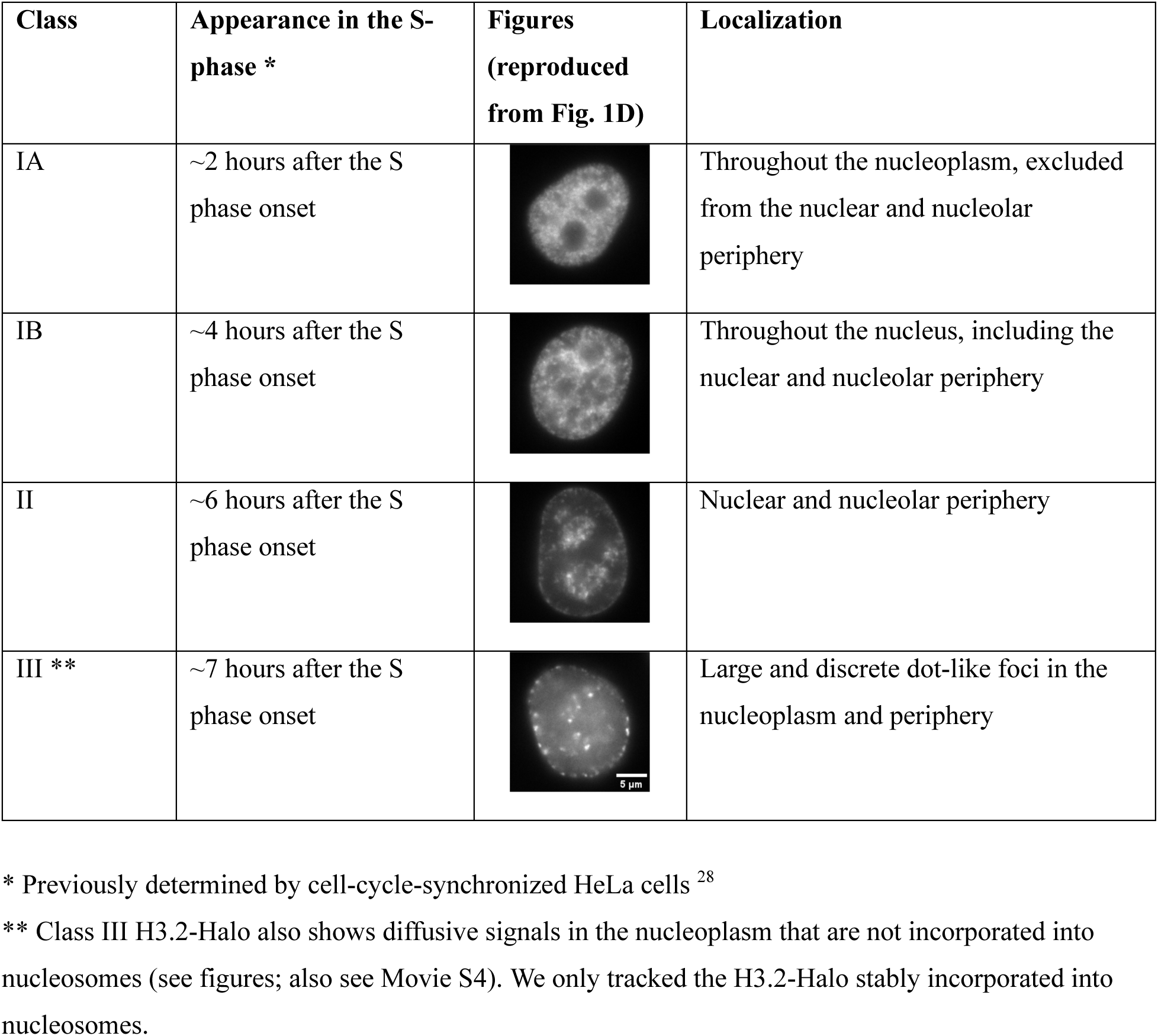
Labeling pattern of the Repli-Histo labeling.

| Class | Appearance in the S-phase * | Figures (reproduced from Fig. 1D) | Localization |
| --- | --- | --- | --- |
| IA | ~2 hours after the S phase onset |  | Throughout the nucleoplasm, excluded from the nuclear and nucleolar periphery |
| IB | ~4 hours after the S phase onset |  | Throughout the nucleus, including the nuclear and nucleolar periphery |
| II | ~6 hours after the S phase onset |  | Nuclear and nucleolar periphery |
| III ** | ~7 hours after the S phase onset |  | Large and discrete dot-like foci in the nucleoplasm and periphery |

\* Previously determined by cell-cycle-synchronized HeLa cells <sup>28</sup>
\*\* Class III H3.2-Halo also shows diffusive signals in the nucleoplasm that are not incorporated into nucleosomes (see figures; also see Movie S4). We only tracked the H3.2-Halo stably incorporated into nucleosomes.

